# Soil organic carbon stocks and stabilization mechanisms in tidal marshes along estuarine gradients

**DOI:** 10.1101/2024.05.18.594814

**Authors:** Friederike Neiske, Maria Seedtke, Annette Eschenbach, Monica Wilson, Kai Jensen, Joscha N. Becker

## Abstract

Tidal marshes store large amounts of soil organic carbon (SOC), however, little is known on SOC stabilization mechanisms in these ecosystems. In estuarine marshes, SOC storage is dominated by a complex interaction of tidal inundation and salinity with biotic ecosystem components, leading to strong spatio-temporal variations within estuaries. Our aim was to assess (i) SOC stocks, (ii) SOC stabilization mechanisms (aggregation and mineral-association), and (iii) their environmental drivers along estuarine gradients. We analyzed SOC stocks and SOC density fractions in topsoil (0-10 cm) and subsoil (10-30 cm) of three marsh zones representing three flooding regimes (daily, monthly, yearly) in three marsh types along the salinity gradient (salt, brackish, freshwater) of the Elbe Estuary, Germany.

Increasing salinity and flooding reduced SOC stocks 0-30 cm (9.3-74.6 t ha^-1^), which was related to decreasing plant biomass and soil texture. Mineral-associated organic matter (C_MAOM_) was the largest SOC fraction (59% of total SOC), followed by aggregate-occluded organic matter (C_oPOM_) (24%) and free particulate organic matter (C_fPOM_) (16%). The C_MAOM_ amount in topsoils decreased downstream with increasing salinity, reflecting decreasing fine-texture along the estuary. The amount of C_oPOM_ was higher in topsoils and high marshes, indicating negative effects of flooding on aggregation. The relative proportion of C_fPOM_ (% of total SOC) increased with increasing flooding frequency and reducing soil conditions.

Our results underline the importance of estuarine gradients as drivers of SOC storage and stabilization. Climate-change induced sea-level rise and variations in salinity might reduce SOC storage and stabilization in estuaries.

## 1 Introduction

Intertidal wetlands play a critical role as carbon (C) sinks on a global scale (Mcleod et al., 2011). They are known for their exceptional capacity to sequester and store significant amounts of soil organic carbon (SOC) (Mcleod et al., 2011; Chmura et al., 2003; Macreadie et al., 2019). Such tidal marshes form the interface of marine and terrestrial ecosystems and are thus shaped by strong environmental gradients. These gradients are particularly pronounced for estuarine marshes with intersecting limnic and marine influence. Increasing salinity towards the coast results in an estuarine salinity gradient forming distinct marsh types along the course of the estuary (salt marshes, brackish marshes, freshwater marshes). Additionally, the elevation of the marsh surface rises from the river inland, leading to a flooding gradient. Both gradients are also reflected in vegetation composition in estuaries (Engels and Jensen, 2009). The interplay of abiotic gradients and biotic communities determine the biogeochemical control of SOC storage in tidal marshes (Seyfferth et al., 2020; Schulte Ostermann et al., 2021) and might be affected by environmental changes (Ruiz-Fernández et al., 2018; Barry et al., 2023; Tang et al., 2023).

Due to their coastal location, tidal marshes and their SOC storage potential are highly vulnerable to climate change (Macreadie et al., 2019; Lovelock and Reef, 2020). In estuaries, a rising sea-level can increase overall saline conditions and lead to the expansion of salt marshes and the displacement of brackish or freshwater marshes (Craft et al., 2009; Visser et al., 2013). Furthermore, a rising sea-level along with an altered flooding regime might force wetlands to move further inland, which is often impeded due to human activities such as embanking (Kirwan and Megonigal, 2013; Enwright et al., 2016). These shifts can have a negative effect on SOC stocks, as previous research indicates that SOC contents decrease with increasing salinity and flooding frequency (Craft, 2007; Spohn et al., 2013; van de Broek et al., 2016). However, previous studies on SOC storage in tidal marshes were mainly conducted in salt marshes (e.g.: Drake et al., 2015; Yuan et al., 2020; Human et al., 2022), while much less is known about tidal freshwater marshes and even less about brackish marshes (e.g.: Spohn and Giani, 2012; van de Broek et al., 2016; Hansen et al., 2017). Considering the pronounced vulnerability of these ecosystems, we need to better understand C dynamics along estuarine gradients and the underlying processes of SOC storage.

In general, SOC storage is regulated by the relation of C inputs and outputs (Lützow et al., 2006). This is particularly important for tidal marshes, which receive large amounts of allochthonous C through flooding-induced deposition and autochthonous OC input through high primary productivity (Mcleod et al., 2011). In contrast, atmospheric CO_2_ outputs are typically low due to limited microbial C mineralization, resulting from anaerobic conditions in regularly flooded soils (Kirwan and Megonigal, 2013; Chapman et al., 2019). This circumstance, together with the recalcitrance of organic matter (OM) itself, has long been considered as the measures for OC protection in wetland soils. Little attention has been paid to alternative mechanisms protecting OC through the interaction with other soil components (van de Broek et al., 2018; Sun et al., 2019; Maietta et al., 2019; Ran et al., 2021). For terrestrial soils, it is well known that the occlusion of OM in aggregates or the association of OM with mineral surfaces (MAOM) increases the accumulation and preservation of OM (Six et al., 2002a; Lützow et al., 2006; Wiesmeier et al., 2019). There is rising evidence that in terrestrial soils, pedogenic stabilization mechanisms are more important for the long-term storage of OC than the recalcitrance of the OM itself. Mineral-associated OM is older, has longer turnover times, and is less responsive to environmental changes than free particulate organic matter (fPOM) (Lützow et al., 2006; Rocci et al., 2021).

For intertidal wetlands, knowledge on effects of mineral association and soil aggregation for SOC stabilization is surprisingly limited and previous studies have reported contradictory results. Maietta et al. (2019) showed that aggregation contributed to stabilization of SOC, while organo-metal oxide complexes were negligible in tidal freshwater marshes of the Patuxent River (Maryland, USA). In contrast, Cui et al. (2014) found that aggregation played a minor role in soils of the subtropical Chongming Island (Yangtze Estuary, China) and the majority of SOC was bound to iron and aluminum oxyhydrates, as well as silt and clay minerals. A reason for these confounding findings may be the high spatio-temporal heterogeneity of biogeochemical properties in estuarine marshes, resulting from strong environmental gradients in salinity and flooding (van de Broek et al., 2016; Seyfferth et al., 2020). The dynamic flooding regime leads to fluctuating redox conditions and can potentially destabilize SOC by the reductive dissolution of iron oxides (Kleber et al., 2015; Chen et al., 2020). Moreover, dry-wet cycles have a strong but opposing effect on aggregate formation and stability which is strongly dependent on soil properties, management practices, or intensities of the dry-wet periods (Six et al., 2004). Dry-wet cycles may increase OM turnover due to the increasing formation of macroaggregates at the expense of more stable microaggregates (Najera et al., 2020). Moreover, inundation may destroy aggregates in soils of regularly flooded ecosystems (Mao et al., 2018; Liu et al., 2021; Ran et al., 2021).

The objectives of our study were to (i) quantify OC stocks along a salinity and flooding gradient of the Elbe Estuary, (ii) estimate SOC stabilization mechanisms (aggregate-occluded and mineral-associated OC), and (iii) relate the SOC storage to site characteristics. Therefore, we quantified SOC stocks and applied a density fractionation to separate OC associated with minerals (C_MAOM_), OC occluded in aggregates (C_oPOM_), and OC in free particulate OM (C_fPOM_). We related our findings to soil conditions (texture, pH, redox conditions) and vegetation properties (aboveground biomass) to identify drivers for SOC storage and SOC stabilization.

## 2 Methods

### 2.1 Study Area

The study was conducted in the Elbe Estuary, Northern Germany (Figure 1). The region is characterized by an oceanic climate with a mean annual temperature of 9.3 °C and mean annual rainfall of 812.8 mm (1991 - 2020, federal state of Schleswig-Holstein) (DWD, 2024). Upstream tidal influence of the Elbe Estuary is restricted by a weir at Geesthacht approximately 140 km inland from the estuaries mouth at Cuxhaven (Boehlich and Strotmann, 2008). The estuary experiences a semi-diurnal meso-to macrotidal regime (Carstens et al., 2004; Boehlich and Strotmann, 2008) with a mean tidal range between 2 m (weir at Geesthacht) and 3.5 m (port of Hamburg) (Kappenberg and Grabemann, 2001). The Elbe Estuary can be subdivided into different salinity zones forming distinct marsh types along the salinity gradient of the estuary (Engels and Jensen, 2009).

**Figure 1:**
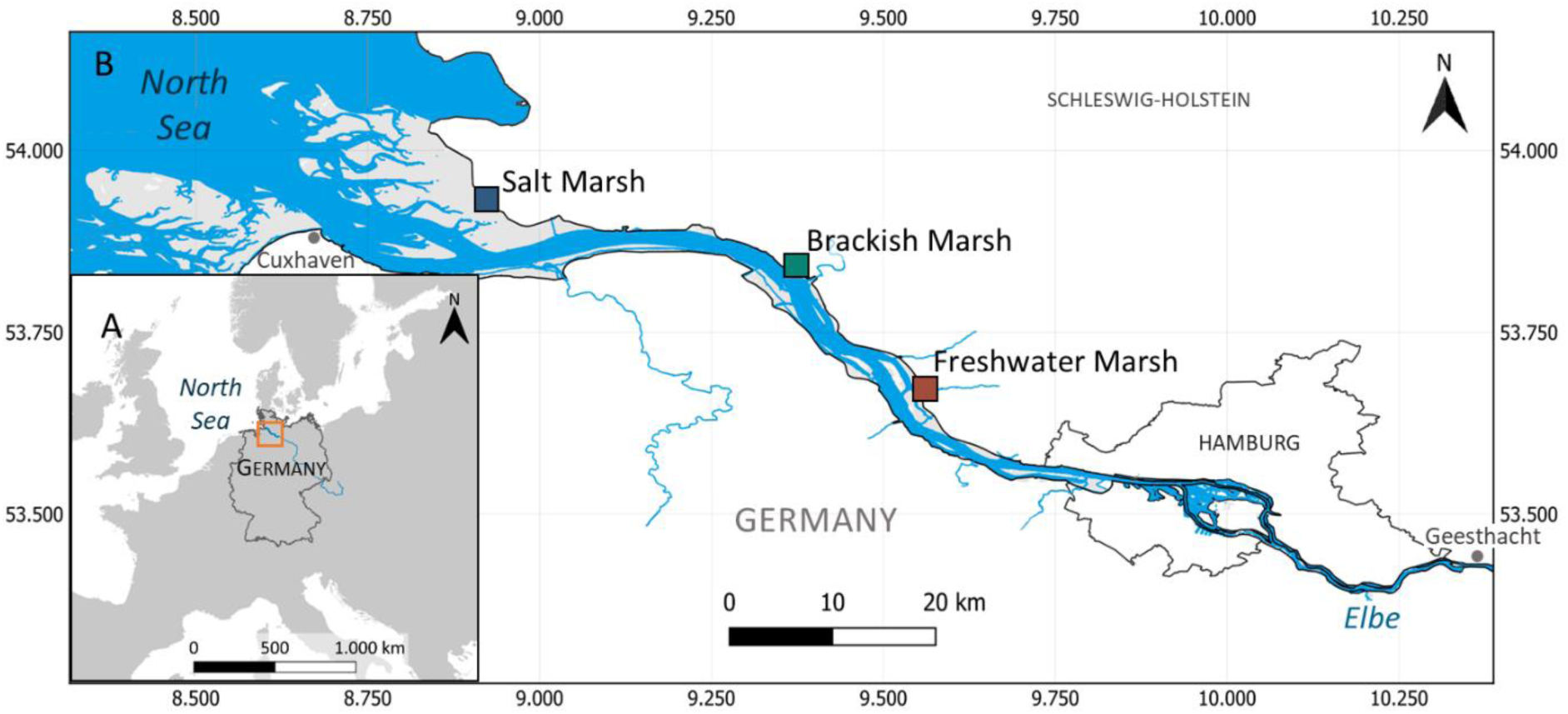
(A) Location of the Elbe Estuary in central Europe (orange rectangle) and (B) location of the marsh sites along the Elbe Estuary (salt marsh: 53°55’40.0’’N 8°54’51.3’’E/river-km: 710; brackish marsh: 53°50’03.0’’N 9°22’15.6’’E/river-km: 680; freshwater marsh: 53°39’56.3’’N 9°33’11.5’’E/river-km: 658) (Data sources: WSV, 2011, 2017; EEA, 2017).

We selected three marsh types along the salinity gradient of the Elbe Estuary (salt marsh, brackish marsh, freshwater marsh). Kaiser-Wilhelm-Koog represents a salt marsh and is located in the Schleswig-Holstein Wadden Sea National Park at the mouth of the estuary. As brackish marsh, a site close to Hollerwettern was chosen and the selected freshwater marsh is located in the nature conservation area “Haseldorfer Binnenelbe” closest to the city of Hamburg. Within each of these marsh types, we selected three locations along the flooding gradient: Pioneer zones (PZ) are located closest to the main channel and experience flooding during high tide (twice per day), low marshes (LM) are flooded during spring tides with new and full moon (occurring twice per month), and high marshes (HM) undergo few inundations per year during storm tides. The different marsh zones and types were defined by dominant plant species that are typical for the respective salinity and flooding regime (Engels and Jensen, 2009) (Table S 1). All locations are situated on the seaward side of a dike and exhibit near-natural conditions.

### 2.2 Field sampling and site characterization

#### Sampling and field site monitoring

Soil samples were collected in February and March 2022 from each marsh type along the salinity gradient (salt marsh, brackish marsh, freshwater marsh) and marsh zone along the flooding gradient (pioneer zone, low marsh, high marsh). Soil samples were taken from five adjacent replicate plots (2 x 2 m) and directly separated into topsoil (0 – 10 cm) and subsoil (10 – 30 cm) samples. At each marsh location, a pipe well was installed for continuous monitoring of (ground-)water level, temperature and salinity (CTD-Diver, vanEssen Instruments, Delft, The Netherlands). Additionally, reference soil profiles were characterized at each site according to WRB (2022) between December 2021 and February 2022. Undisturbed soil cores (volume: 100 cm^3^) for bulk density analysis were taken from each soil profile (n = 5) at depths of 5, 10, 30, and 60 cm. Aboveground plant biomass was harvested from two 20 x 20 cm squares in each of the five replicate plots in late July 2022.

#### Soil Reduction Index

Reducing soil conditions at each replicate plot were investigated using the “Indicator of Reduction in Soil” (IRIS) (Castenson and Rabenhorst, 2006; Rabenhorst, 2008). In this study, the method was modified based on Mueller et al. (2020) and Mittmann-Goetsch et al. (2024, Preprint). In short, PVC sticks (5 x 70 cm) were covered with FeCl_3_-paint and inserted into the soil down to a depth of 60 cm for four weeks. After retrieval of the sticks, a reduction index (RI) (0 – 1) was calculated based on the area where the FeCl_3_-paint was removed from the PVC sticks. Field incubation of IRIS sticks were conducted over the course of almost one year (11 x 4 weeks). We calculated the reduction index over the whole period (RI_year_) and for the soil sampling period in February and March (RI_march_) for depth increments corresponding to the soil sampling depths (0 – 10 cm, 10 – 30 cm).

### 2.3 Laboratory analyses

Laboratory analyses were conducted at the Institute of Soil Science, University of Hamburg unless indicated otherwise. Soil samples were air-dried at room temperature until constant weight and sieved to 2 mm.

#### Density fractionation

A density fractionation was applied on the soil samples to separate the SOC into the free light fraction (which corresponds to free particulate organic matter = fPOM), occluded light fraction (organic matter occluded in aggregates = oPOM), and heavy fraction (mineral-associated organic matter = MAOM) based on Golchin et al. (1994) and Viret and Grand (2019). A sodium polytungstate (SPT) (3 Na_2_WO_4_ · 9 WO_3_ · H_2_O) solution (ρ = 1.62 g cm^-3^) was added to the air-dried and sieved soil samples. The suspensions were carefully mixed and, after resting for 30 minutes, centrifuged at 3000 rpm for 60 minutes. The supernatants were filtered (cellulose acetate filter, 0.45 μm) under vacuum. The filtrates, consisting of fPOM, were washed with distilled water to remove the SPT solution until the supernatants reached an electrical conductivity (EC) of < 50 μS cm^-1^. Afterwards, the fPOM was dried at 65 °C. In order to separate the oPOM from the remaining soil deposits, the sample was mixed again with SPT solution and treated with ultrasound (171 W ml^-1^) for 11 minutes to disrupt soil aggregates. The suspended samples were centrifuged at 3000 rpm for 60 minutes, supernatants (consisting of oPOM) were vacuum filtered, and filtrates were washed with distilled water and dried at 65 °C. The remaining soil (containing MAOM) was washed with distilled water and centrifuged (3000 rpm for 60 minutes) until the supernatant’s EC was < 50 μS cm^-1^. The obtained MAOM was dried at 105 °C until constant weight.

#### Determination of C and N contents (soil)

The MAOM and bulk soil samples were ground and dried at 105 °C, before they were analyzed for their inorganic and organic C contents with an elemental analyzer (soli TOC® cube, Elementar Analysensysteme GmbH, Langenselbold, Germany) and total C and N contents (vario Max cube, Elementar Analysensysteme GmbH, Langenselbold, Germany). Samples of fPOM and oPOM were ground and as sample volumes were very low, OC and N contents were analyzed during isotope ratio mass spectrometry (Delta XP, Thermo Electron, Bremen Germany; with a preconnected elemental analyzer Flash EA 1112, Thermo Electron, Rodano, Milano, Italy) at the Centre for Stable Isotope Research and Analysis (KOSI) at the University of Goettingen. Organic carbon associated with each fraction (C_MAOM,_ C_oPOM,_ C_fPOM_) was expressed as total amounts (mg C g^-1^ soil) or as relative proportions of total SOC (% of total SOC).

#### Calculation of C Stocks

Soil organic carbon stocks (t ha^-1^) were calculated with the following equation:

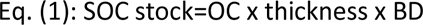

where OC is the OC content (%), thickness is the thickness of the soil layer (cm) and BD is the bulk density of the soil layer (g soil cm^-3^).

#### General soil characterization

Soil texture was determined using the sieving and sedimentation method for mineral soils (Müller et al., 2009). Soil samples were freed from organic matter (samples with OC content > 1%) and carbonates (samples with carbonate content > 0.4%) by treatment with 30% H_2_O_2_ and hydrochloric acid (10% HCl solution), respectively. After dispersion with sodium pyrophosphate (Na_4_P_2_O_7_ **·** 10 (H_2_O)), the sand fraction (63 μm - 2000 μm) was assessed with a vibratory sieve shaker (Retsch GmbH, Haan, Germany). The fine fraction was analyzed based on the Köhn-pipette fractionation method in a Sedimat 4–12 (Umwelt-Geräte-Technik GmbH, Muencheberg, Germany). Soil pH was measured with a pH meter (MP230 GLP, Mettler-Toledo GmbH, Gießen, Germany) in a 0.01 M CaCl_2_ solution (pH_CaCl2_) and in H_2_O (pH_H2O_). The soil-water-suspension was used to determine the EC with a conductivity meter (WTW Cond 330i with TetraCon 325, Xylem Analytics Germany Sales GmbH & Co. KG, Weilheim, Germany).

#### Aboveground plant biomass

Plant samples were washed with water to remove adhered sediments and dried at 60 °C for at least 48 h until constant weight. After drying, plant samples were weighed to estimate the dry weight. Dried plant samples were ground and analyzed for total C and total N content (vario Max cube, Elementar Analysensysteme GmbH, Langenselbold, Germany).

### 2.4 Data analysis

Differences between marsh locations were assessed by analysis of variance (ANOVA) followed by TukeyHSD post-hoc comparison. Depth effects were assessed separately for each marsh location using a linear mixed effect model (LME) for paired comparison with depth as a main factor and sampling location as random factor. Assumptions of normality and variance homogeneity were checked by visual inspection of model residuals. When necessary, Dixon’s Q test was applied to check groups for potential outliers. Relationships between measurement variables were assessed by linear regression and Pearson correlation. Additionally, Spearman rank correlation coefficients were included to account for potential non-linearity. Statistical differences were accepted as significant at p-level < 0.05 and p-levels between 0.10 and 0.05 were considered as significance by tendency. Statistical analyses were conducted in R 4.2.0 (R Core Team, 2022), using “multcomp” (Hothorn et al., 2016) and “multcompView” (Graves et al., 2015) packages, as well as “ggplot2” (Wickham et al., 2016) for data visualization.

## 3 Results

### 3.1 Effect of estuarine gradients on SOC storage

#### Soil organic carbon storage along gradients

Soil organic carbon storage (SOC stocks and contents) were affected by the estuarine gradients and depth (Figure 2, Table S 2 + 3). The SOC stocks in 0 - 30 cm depth ranged from 9.3 to 74.6 t ha^-1^ across the examined marshes in the Elbe Estuary and were partly negatively affected by increasing salinity and flooding (Figure 2). The highest SOC stocks were found in the freshwater marsh (69.2 - 74.6 t ha^-1^) and in all high marshes (67.6 - 73.7 t ha^-1^), while the brackish pioneer zone had the lowest SOC stocks (9.3 ± 1.4 t ha^-1^). The SOC stocks in the pioneer zones and low marshes decreased by 33% and 46% from freshwater to salt marsh conditions, respectively, while the high marshes were not affected by the salinity gradient. The SOC stocks generally decreased with more frequent flooding, but the decrease was absent in the freshwater marsh. The strongest decline along the flooding gradient occurred in the brackish marsh, with a reduction of 86% from the high marsh to pioneer zone. The SOC contents in bulk soil varied between 0.23% and 5.8% in the different marshes and depths (Table S 2). The SOC contents decreased with increasing salinity (except for high marsh subsoils) and with increasing flooding at the salt and brackish marsh (Table S 3). At the freshwater marsh, the SOC contents did not change along the flooding gradient. Effects of the estuarine gradients on SOC contents were more pronounced in topsoils compared to subsoils.

**Figure 2:**
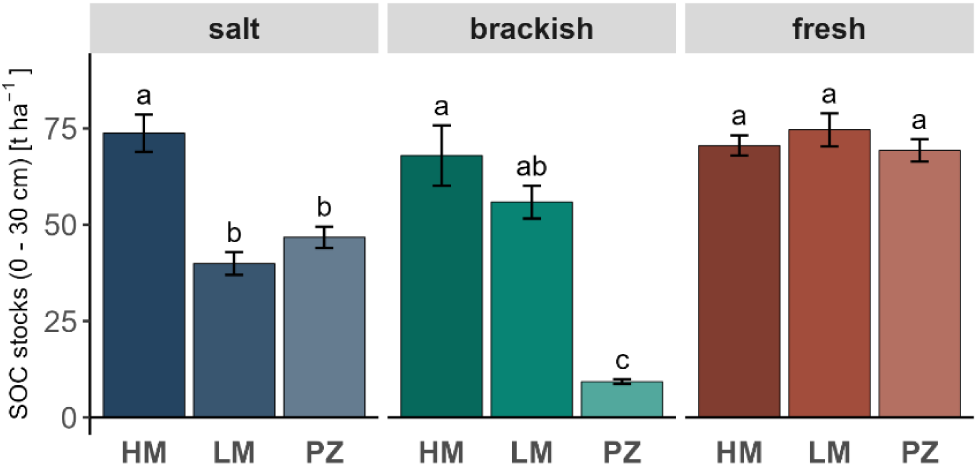
Soil organic carbon (SOC) stocks [t ha^-1^] (0 – 30 cm) of the different marsh types and zones along the salinity (salt marsh, brackish marsh, freshwater marsh) and flooding gradient (HM = high marsh, LM = low marsh, PZ = pioneer zone) of the Elbe Estuary. Error bars indicate standard error of the mean (n = 5). Lowercase letters indicate significant differences (p < 0.05) derived from ANOVA with TukeyHSD post-hoc comparison.

### 3.2 Effect of estuarine gradients on total amounts of SOC fractions

Mineral-associated OC (C_MAOM_) made up the largest SOC fraction in terms of total amount in the investigated marshes with an average amount of 15.4 ± 11.7 mg C g^-1^ (Figure 3 A). The amount of C_MAOM_ decreased with increasing salinity with a more pronounced effect in topsoils and varied along the flooding gradient with no consistent trend. In topsoils, high amounts of C_MAOM_ were found in the freshwater marsh (28.7 ± 10.4 mg C g^-1^) and decreased by −56% towards the salt marsh with the strongest decline in pioneer zones (−69%). In subsoils, the C_MAOM_ amount significantly decreased with increasing salinity only in the pioneer zones (−71%). Overall, the brackish pioneer zones had the lowest amounts of C_MAOM_ with 0.9 ± 0.2 mg C g^-1^ in topsoil and 1.3 ± 0.5 mg C g^-1^ in subsoil. The amount of C_MAOM_ decreased along the flooding gradient from the high marsh to the pioneer zone in the brackish topsoil (−96%) and subsoil (−83%), as well as in the subsoil of the salt marsh (−40%).

**Figure 3:**
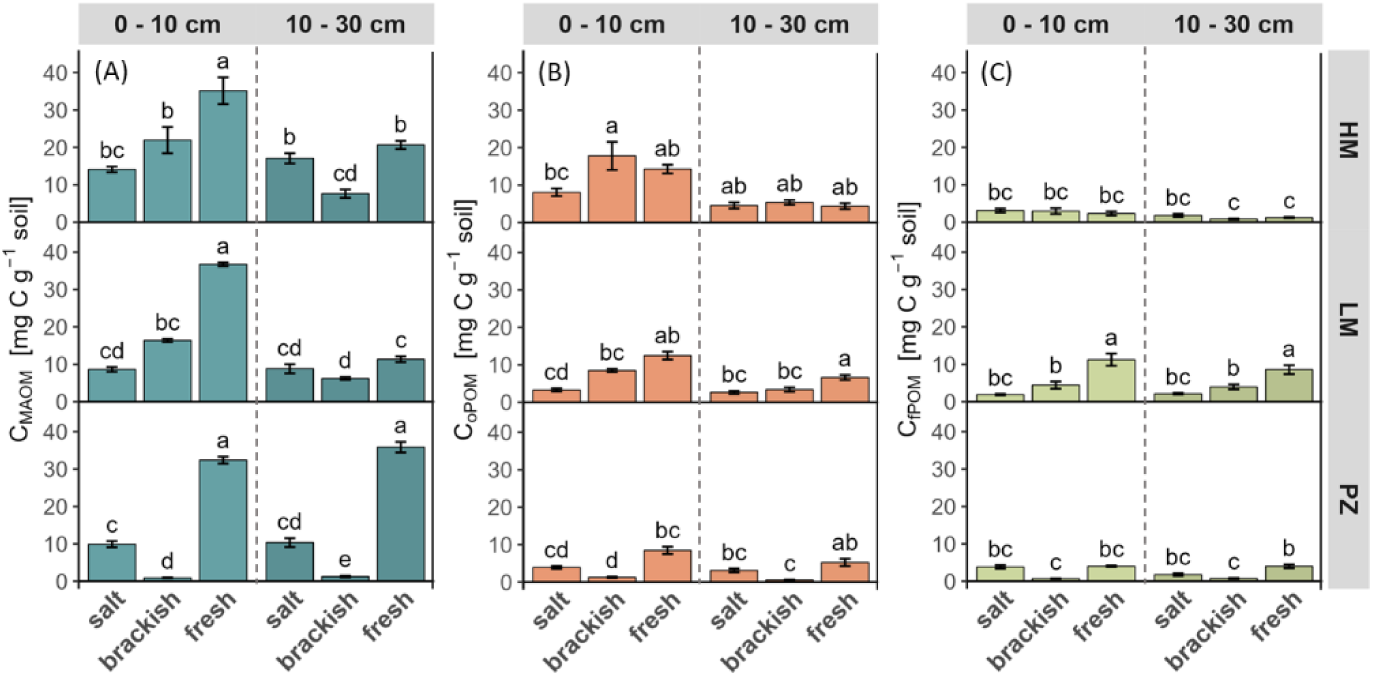
Amount (mg C g^-1^ soil) of mineral-associated OC (C_MAOM_) (A), aggregate-occluded OC (C_oPOM_) (B), and OC in free particulate organic matter (C_fPOM_) (C) of the different marsh types and zones along the salinity (salt marsh, brackish marsh, freshwater marsh) and flooding gradient (HM = high marsh, LM = low marsh, PZ = pioneer zone) of the Elbe Estuary. Error bars indicate standard error of the mean (n = 5). Lowercase letters indicate significant differences (p < 0.05) among marsh types and zones per SOC fraction and per depth derived from ANOVA with TukeyHSD post-hoc comparison.

Occluded POM in aggregates (C_oPOM_) represents the second largest SOC fraction with 5.6 ± 4.6 mg C g^-1^ (Figure 3 B). The estuarine gradients had no clear effect on the amount of C_oPOM_ but in some marshes the C_oPOM_ amount showed a negative response to increasing salinity and flooding frequency. The highest amounts of C_oPOM_ were found in topsoils of the brackish high marsh (17.8 ± 8.4 mg C g^-1^), freshwater high marsh (14.3 ± 2.6 mg C g^-1^) and freshwater low marsh (12.5 ± 2.4 mg C g^-1^), while the lowest amounts were detected in subsoil of the brackish pioneer zone (0.5 ± 0.3 mg C g^-1^). The amount of C_oPOM_ showed a significant decrease along the salinity gradient from the freshwater to the salt marsh in low marsh topsoils (−73%) and subsoils (−60%). The flooding gradient also tended to have a negative effect but this was only significant in the brackish marsh with a reduction from high marshes to pioneer zones by −93% in topsoils and −90% in subsoils. Overall, topsoils (8.7 ± 5.4 mg C g^-1^) had more than twice the C_oPOM_ amount of subsoils (4.0 ± 1.8 mg C g^-1^) but a significant decrease with increasing depth was only observed in parts of the brackish and freshwater marsh.

Free POM was the smallest SOC fraction with 3.17 ± 2.7 mg C g^-1^ (Figure 3 C). The amount of C_fPOM_ decreased with increasing salinity in low marshes while the flooding gradient did not exhibit a clear effect. Highest amounts of C_fPOM_ were found in topsoil (11.2 ± 3.6 mg C g^-1^) and subsoil (8.6 ± 2.6 mg C g^-1^) of the freshwater low marsh, from where it decreased along the salinity gradient towards the salt marsh by −83% (topsoil) and −75% (subsoils). The flooding gradient did not exhibit a clear trend on the amount of C_fPOM_: The amount of C_fPOM_ remained relatively constant along the flooding gradient at the salt marsh while it decreased in the brackish topsoil by −93% from the high marsh to the pioneer zone. In contrast, the amount of C_fPOM_ was positively influenced by an increasing flooding in the freshwater subsoil (+16%).

### 3.3 Effect of estuarine gradients on relative proportions of SOC fractions

Proportions of C_MAOM_ constituted the largest proportion of total SOC (59.2 ± 11.2% of total SOC) ranging between 42% and 82% of total SOC (Figure 4 A). Proportions of C_MAOM_ decreased with increasing salinity in topsoils while the flooding did not exhibit an overall trend on C_MAOM_ proportions. Maximum proportions of C_MAOM_ were found in freshwater soils (67.3 ± 11.2%). Topsoil proportions of C_MAOM_ decreased from the freshwater to the salt marsh along the salinity gradient by −27% in high marshes and −22% in pioneer zones. The brackish marsh tended to have lower C_MAOM_ proportions (49.9 ± 11.2% in topsoil and subsoil combined) compared to salt and freshwater marshes.

**Figure 4:**
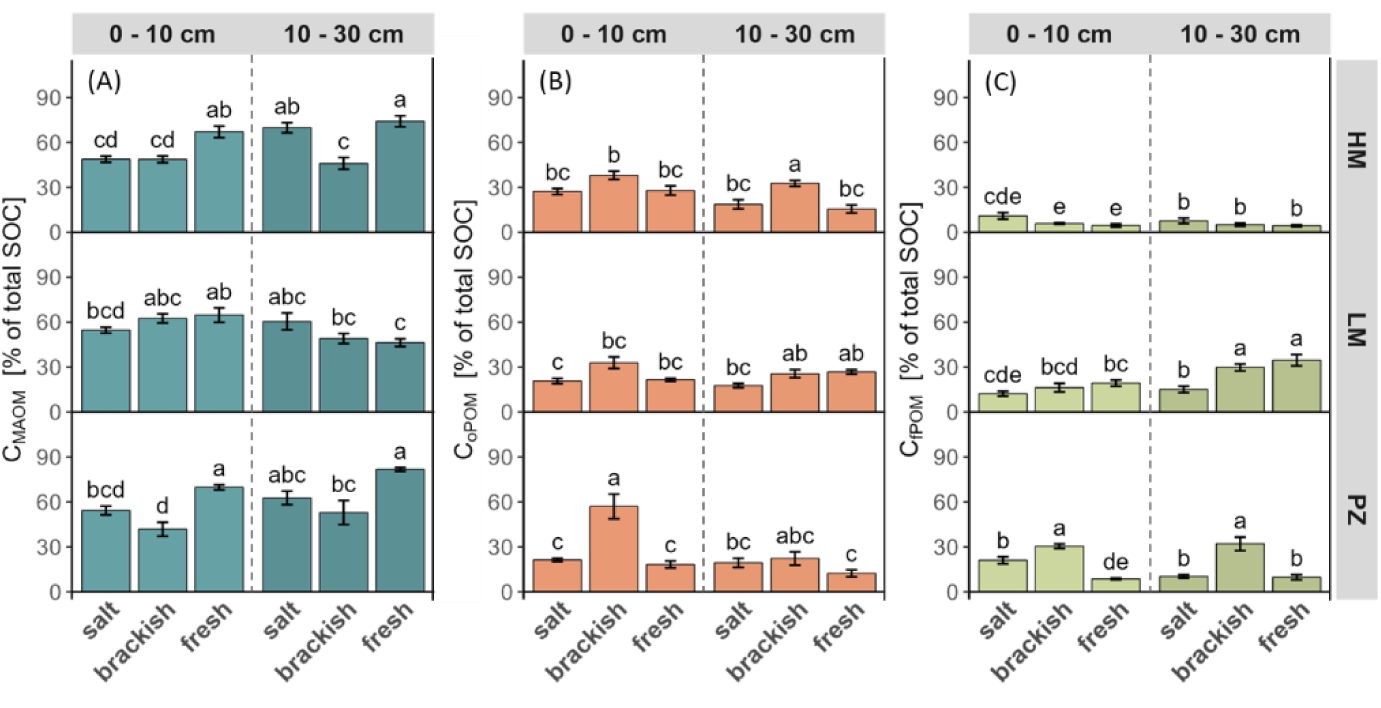
Relative proportions of mineral-associated OC (C_MAOM_) (A), aggregate-occluded OC (C_oPOM_) (B), and OC in free particulate organic matter (C_fPOM_) (C) of total SOC (% of total SOC) of the different marsh types and zones along the salinity (salt marsh, brackish marsh, freshwater marsh) and flooding gradient (HM = high marsh, LM = low marsh, PZ = pioneer zone) of the Elbe Estuary. Error bars indicate standard error of the mean (n = 5). Lowercase letters indicate significant differences (p < 0.05) among marsh types and zones per SOC fraction and per depth derived from ANOVA with TukeyHSD post-hoc comparison.

Occluded POM presented the second largest proportion of total SOC (24.2 ± 10.3%) with the highest values in the brackish marsh (Figure 4 B). Proportions of C_oPOM_ did not show a uniform trend along the estuarine gradients. The brackish marsh exhibited the highest C_oPOM_ proportions (32.1 ± 10.3%) with maximum values in the topsoil of the brackish pioneer zone (57.0 ± 18.4%) while lowest proportions were found in subsoil of the freshwater pioneer zone (12.4 ± 5.1%). Overall, proportions of C_oPOM_ decreased from top-to subsoil (29.5 ± 12.2% to 21.2 ± 6.3%).

Free POM had the lowest proportion of SOC (15.8 ± 10.2%) (Figure 4 B). The salinity gradient did not display a uniform effect, while increasing flooding frequency exhibited a positive effect on C_fPOM_ proportions. The highest proportions of C_fPOM_ were found in subsoils of the freshwater low marsh (34.6 ± 8.4%), brackish low marsh (29.8 ± 5.2%), and in the brackish pioneer zone (30.5 ± 3.4% - 32.1 ± 10.2%). While C_fPOM_ proportions in most marshes remained largely unaffected by the salinity gradient, the proportions doubled along the salinity gradient from the freshwater to the salt marsh in topsoils of pioneer zones and were reduced by 50% in the subsoils of low marshes. The lowest proportions of C_fPOM_ occurred in high marshes (6.2 ± 2.5%) and increased along the flooding gradient towards pioneer zones (18.3 ± 10.7%). The strongest increase along the flooding gradient occurred at the brackish marsh, where C_fPOM_ proportions were five (topsoils) to six times (subsoils) higher in the pioneer zone compared to the high marsh. Proportions of C_fPOM_ also nearly doubled along the flooding gradient from the high marsh to the pioneer zone in topsoil of the salt marsh.

### 3.4 Soil properties

The soil texture was affected by estuarine gradients but trends were not uniform along the estuary (Table 1). The highest clay contents were found in the freshwater low marsh (topsoil: 39.8%) and freshwater pioneer zone (topsoil: 39.9%; subsoil: 37.0%). The brackish marsh tended to have lower clay (3.3% - 24.4%) and silt (11.8% - 63.4%) but higher sand contents (12.2% - 82.0%). The EC increased along the salinity gradient from the freshwater marsh (104 - 1119 µS cm^-1^) to the salt marsh (2080 - 3134 µS cm^-1^). Pronounced differences in redox conditions (RI_year_ and RI_march_) were observed with respect to soil depth and the estuarine gradients. The average RI_year_ over the year (0.2) decreased downstream with increasing salinity in pioneer zones. Highest values of RI_year_ were found in subsoil of pioneer zones (0.23 - 0.66), while topsoils of low and high marshes exhibited the lowest RI_year_ (0.03 - 0.09). The average RI for the soil sampling period (RI_march_) was 0.2 and showed a less pronounced trend than RI_year_.

**Table 1:**
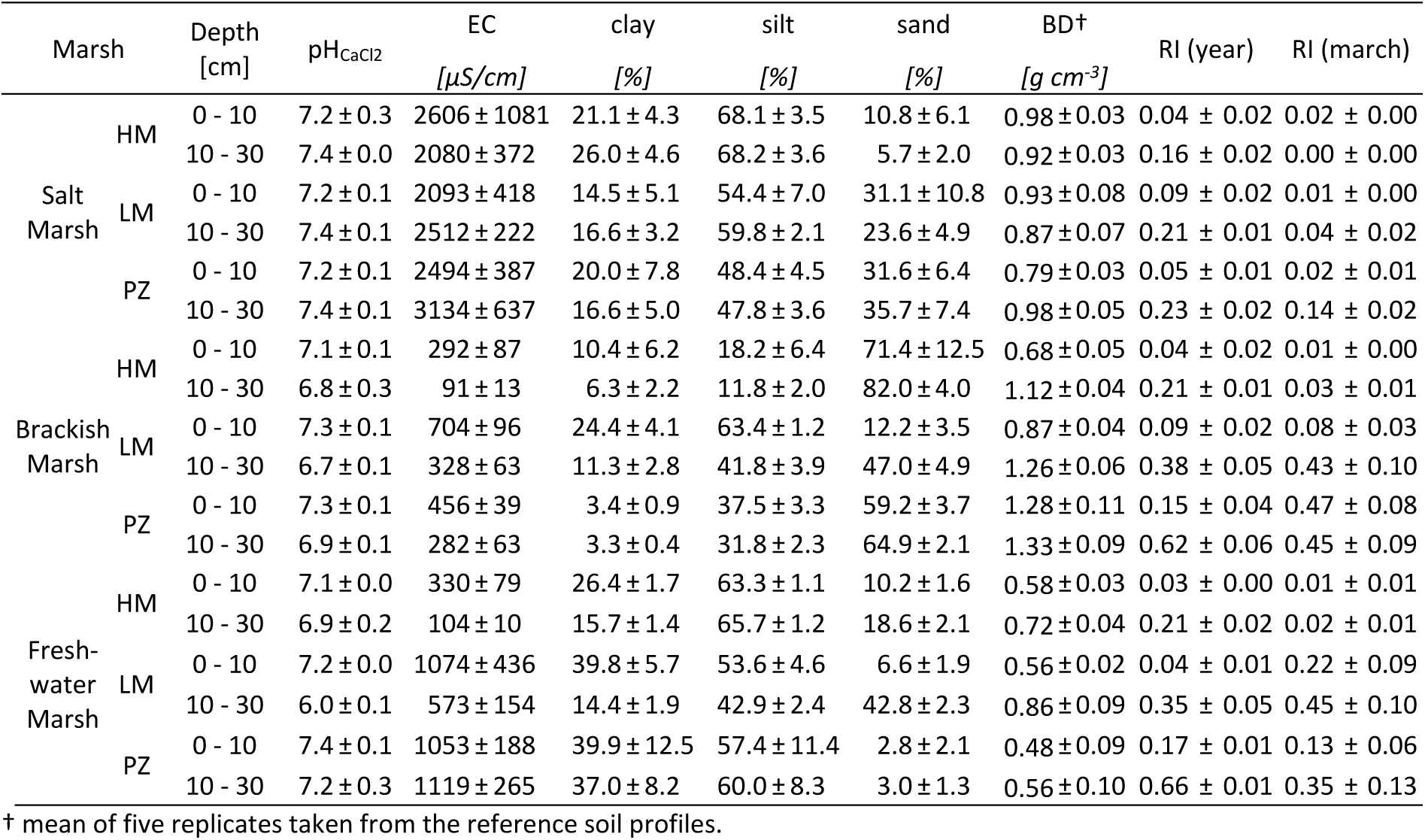
Soil characteristics (*pH_CaCl2_* = soil pH measured in CaCl_2_-solution; *EC* = soil electric conductivity; *BD* = soil bulk density; *RI (year)* = mean soil reduction index of one year derived from IRIS method; *RI (march)* = soil reduction index for the sampling period in February and March derived from the IRIS method) of the different marsh types and zones along the salinity (salt marsh, brackish marsh, freshwater marsh) and flooding gradient (HM = high marsh, LM = low marsh, PZ = pioneer zone) of the Elbe Estuary (mean ± standard error, n = 5).

### 3.5 Aboveground plant biomass

The highest aboveground plant biomass was found in the freshwater marsh and the brackish high marsh (Table 2). The aboveground plant biomass decreased downstream with increasing salinity in low marshes (−75%) and pioneer zones (−62%). No clear trends in the C:N ratio of the aboveground plant biomass along the estuary were observed (Table 1). Ratios decreased downstream with increasing salinity in pioneer zones, while ratios did not change along high and low marshes. A decrease in C:N ratio with increasing flooding frequency was detected at the salt marsh while the C:N ratios increased with increasing flooding frequency at the brackish marsh.

**Table 2:** Total biomass (g DW m^-2^) and C:N ratio of the aboveground vegetation present at the different marsh types and zones along the salinity (salt marsh, brackish marsh, freshwater marsh) and flooding gradient (HM = high marsh, LM = low marsh, PZ = pioneer zone) of the Elbe Estuary (mean ± standard error, n = 5).

| Marsh |  | aboveground vegetation |  |
| --- | --- | --- | --- |
|  |  | biomass<br>[g DW m<sup>-2</sup>] | C:N ratio |
| Salt Marsh | HM | 1427.5 ± 174 | 39.9 ± 4.4 |
|  | LM | 860.4 ± 42 | 38.8 ± 7.1 |
|  | PZ | 960.2 ± 55 | 26.8 ± 3.2 |
| Brackish Marsh | HM | 2421.0 ± 365 | 37.1 ± 4.2 |
|  | LM | 2213.6 ± 314 | 52.2 ± 6.1 |
|  | PZ | 1367.9 ± 140 | 48.3 ± 2.2 |
| Fresh-water Marsh | HM | 1999.0 ± 421 | 43.3 ± 6.9 |
|  | LM | 3404.4 ± 234 | 43.9 ± 2.7 |
|  | PZ | 2504.3 ± 235 | 49.0 ± 5.7 |

### 3.6 Relationships between environmental variables and SOC storage and stabilization

Soil organic carbon storage (SOC stocks and contents) had a strong positive correlation with proportions of C_MAOM_ and the amount of all three SOC fractions (C_MAOM_, C_oPOM_, C_fPOM_) (Figure 5 A + B). The clay content was a good predictor for SOC storage, for the amount of C_MAOM_ in topsoil and subsoil, and the amount of C_oPOM_ in topsoil. Proportions of C_MAOM_ also showed a positive correlation with clay content in topsoil and subsoil. The amounts and proportions of C_oPOM_ were negatively correlated with EC with the exception of the C_oPOM_ amount in subsoil. The SOC storage in topsoils correlated negatively with the RI_year_, while RI_march_ positively affected amounts and proportions of C_fPOM_. Aboveground biomass was positively correlated with SOC storage in topsoil as well as the amount of C_MAOM_ in topsoil, C_oPOM_ in topsoil and subsoil and C_fPOM_ in topsoil and subsoil. The majority of the fraction proportions were unaffected by aboveground plant biomass. The C:N ratio of the aboveground plant biomass had negligible effects on SOC stocks, SOC contents, and SOC fractions.

**Figure 5:**
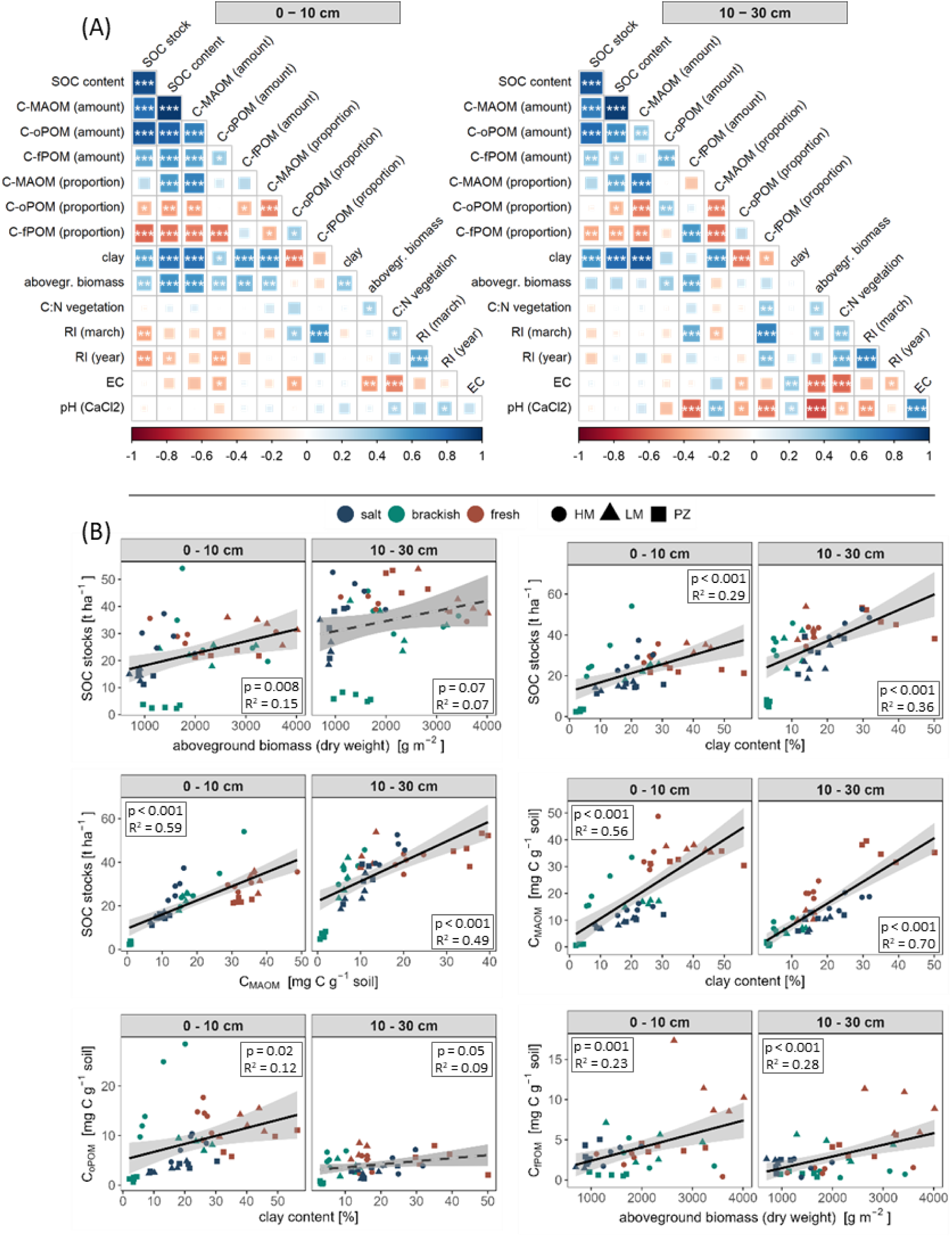
(A) Pearson correlation matrices for topsoil (0 – 10 cm) and subsoil (10 – 30 cm) (n = 45). Asterisk indicate significant correlation at * p < 0.05, ** p < 0.005, and *** p < 0.001. (B) Linear regression between SOC stocks, SOC fractions (C_MAOM_ = mineral-associated OC, C_oPOM_ = OC associated with aggregate-occluded OC, C_fPOM_ = OC in free particulate organic matter), and selected site characteristics (n = 45).

## 4 Discussion

### 4.1 Effect of estuarine gradients on SOC storage

The SOC stocks (0 - 30 cm) in the Elbe Estuary varied between 9.3 – 74.6 t ha^-1^. This is at the lower end of previously described global medians of 79.2 ± 38.1 t ha^-1^ SOC in tidal marshes (0 - 30 cm) (Maxwell et al., 2023). Previous research on North Sea estuaries found SOC stocks between 99 – 217 t ha^-1^ (0 – 60 cm, Scheldt Estuary) (van de Broek et al., 2016), 40 – 134 t ha^-1^ (0 – 80 cm, Elbe Estuary) (Schulte Ostermann et al., 2021), or 31 – 153 t ha^-1^ (0 – 30 cm, Elbe Estuary) (Hansen et al., 2017). The SOC storage (SOC stocks and contents) in our study were predominantly negatively affected by higher salinity and flooding frequency, which is in line with previously reported research from Europe (van de Broek et al., 2016; Hansen et al., 2017; Mazarrasa et al., 2023), temperate USA (Craft, 2007; Kauffman et al., 2020), Australia (Kelleway et al., 2016; Gorham et al., 2021), or China (Yuan et al., 2020). These trends are likely related to differences in OM inputs and preservation capacity along estuarine gradients. As estuarine tidal marshes form the interface of marine, riverine, and terrestrial ecosystems, soils experience a quantitative and qualitative change of autochthonous (local biomass) and allochthonous (flood deposits and sedimentation) OC input along the salinity and flooding gradients (Abril et al., 2002; Hansen et al., 2017). In accordance with the decreasing SOC stocks, autochthonous OM inputs and OC contents in deposited sediments generally decrease from freshwater to salt marshes and with higher flooding frequency (Butzeck et al., 2015). Moreover, higher SOC contents in high marshes might be also attributed to a progressed soil development with continuous C accumulation (Spohn et al., 2013).

The aboveground plant biomass, along with its potential autochthonous OC input, decreased downstream with increasing salinity at the Elbe Estuary and was positively correlated to SOC stocks, as well as SOC contents. Due to the high primary productivity in tidal marshes, the local plant biomass can strongly contribute to SOC storage in marshes (Mcleod et al., 2011; Yuan et al., 2020). The plant biomass production is generally greater at freshwater and brackish marshes than at salt marshes due to reduced salinity stress and higher nutrient availability in freshwater marshes (Więski et al., 2010; Hansen et al., 2017). Consistent with these findings, a higher contribution of tidal marsh vegetation to SOC stocks in fluvial compared to marine settings were found in estuarine marshes of temperate Australia (Gorham et al., 2021). The aboveground biomass of the Elbe Estuary also tended to decrease with increasing flooding frequency at the salt and brackish marsh (p < 0.1), which could have contributed to the observed negative trends of SOC stocks with increasing flooding frequency. However, not all our sites expressed a positive relationship between aboveground plant biomass and SOC storage. Other controlling factors, such as local geomorphology, allochthonous OC inputs, or the OC preservation capacity need to be considered as well (van de Broek et al., 2016; Kelleway et al., 2016; van de Broek et al., 2018). Allochthonous OC inputs can exceed the amount of autochthonous OC input in marshes (Zhou et al., 2007; van de Broek et al., 2016), and can dominate the long-term SOC storage in these soils (van de Broek et al., 2018). The allochthonous OC input into the marsh is determined by the concentration of suspended sediments and associated OM in the river water, both of which show high spatio-temporal variations along the course of estuaries (Abril et al., 2002; Butzeck et al., 2015; Kelleway et al., 2016). It was found that the sediment concentration in the channel of the Elbe Estuary declines downstream (Abril et al., 2002) which might result in a reduced deposition of sediments and accompanied OC into the marsh (Butzeck et al., 2015; van de Broek et al., 2018). Therefore, lower SOC stocks in the salt marsh of the Elbe Estuary are mainly coinciding with low autochthonous and allochthonous OC inputs. Moreover, it was found that freshwater marshes exhibit a greater SOC preservation capacity compared to marine settings due to their fine-grained texture (Kelleway et al., 2016; van de Broek et al., 2018; Gorham et al., 2021). Accordingly, we found larger SOC stocks with higher clay contents. Fine texture can increase SOC storage due to lower drainage and oxygen exchange, leading to lower redox potentials and mineralization rates (Kirwan and Megonigal, 2013; Chapman et al., 2019). However, we found only a weak negative relationship between SOC storage and proxies of inhibited OC mineralization e.g. the reduction index calculated for the whole year (R^2^ < 9% for SOC content and R^2^ < 20% for SOC stocks). Oxygen limitation seems therefore not to explain the observed patterns of SOC storage along the estuarine gradients of the Elbe Estuary. Fine minerogenic particles also have a strong potential to protect OC against mineralization by building organo-mineral interactions or by the occlusion in aggregates (Wiesmeier et al., 2019; Six et al., 2002a; Six et al., 2002b). Therefore, the positive effect of clay on SOC contents might indicate that SOC stabilization mechanisms by mineral association or aggregation play important roles for SOC storage in the Elbe Estuary.

### 4.2 SOC stabilization mechanisms along estuarine gradients

The majority of SOC in the Elbe marshes was associated to minerals (C_MAOM_: 59%), indicating that organo-mineral interaction plays a key role in the Elbe Estuary. Other SOC fractions (C_oPOM_: 24% and C_fPOM_: 16%) were less abundant. This is in contrast to a study in tidal freshwater marshes of the Patuxent River in Maryland (USA) where SOC was predominantly stored in aggregates and not in organo-mineral complexes (Maietta et al., 2019). However, the importance of mineral association for SOC storage in salt marshes was also emphasized (Sun et al., 2019). These contrasting results indicate that SOC stabilization mechanisms undergo strong spatial variability between estuaries as well as along estuarine gradients.

We found highest amounts of C_MAOM_ at the freshwater marsh of the Elbe Estuary and a partially decreasing trend along the salinity gradient from the freshwater to the salt marsh. These trends, for the most part, were positively related to aboveground biomass and clay content. The important role of clay for SOC stabilization is well established due to the provision of surface area for OM adsorption and organo-mineral interactions (Six et al., 2002a; Lützow et al., 2006; Wiesmeier et al., 2019). The contribution of fine minerogenic particles (< 63 μm) and OC to the amount of deposited sediments in the Elbe Estuary is generally higher in the freshwater compared to the salt marsh (Butzeck et al., 2015). This decreasing input of fine particles and OM downstream of the estuary reduces the input of new mineral surface areas for mineral association of OC and might indicate a decreased input of already mineral-associated OC. Therefore, the decreasing trend of C_MAOM_ downstream of the estuary from the freshwater to the salt marsh might be the combined effect of decreasing input of OC (autochthonous and allochthonous) and mineral surfaces by sedimentation along the gradient. This is supported by a maximum occurrence of C_MAOM_ in topsoils and a strongly expressed effect of the salinity gradient in pioneer zones, which are more affected by recent sedimentation. The contribution of C_MAOM_ to total SOC in pioneer zones and high marshes decreased downstream with increasing salinity, while proportions of C_fPOM_ partly increased. This underlines the decreasing formation and relevance of mineral association for SOC storage from the freshwater to the salt marsh of the Elbe Estuary. Similar to C_MAOM_, the amount of C_oPOM_ was also largely related to aboveground biomass and clay content, and showed a decreasing trend from the freshwater to the salt marsh. Fine mineral particles play an import role in the formation of aggregates (Six et al., 2002b; Lützow et al., 2006). Thus, the decreasing trend of C_oPOM_ with increasing salinity might be attributed to the decreased availability of clay for aggregation due to a reduced deposition of fine mineral particles in salt marshes compared to freshwater marshes. These observed negative trends downstream of the estuary in aggregation might be further amplified by the direct adverse effect of salt on aggregate stability (Wong et al., 2010). This is supported by a negative correlation between EC of the marsh soils and the amounts and proportions of C_oPOM_. The high amounts of C_fPOM_ in the freshwater marsh and its decreasing trend downstream of the estuary with increasing salinity is strongly linked to the local aboveground plant biomass. In contrast, the increasing contribution of C_fPOM_ to total SOC downstream with increasing salinity in pioneer zones and high marshes highlights the shift from protected SOC in mineral association in the freshwater marsh towards pedogenically unprotected free POM in the salt marsh.

The flooding gradient had no clear effect on the amount of C_MAOM_ and C_oPOM_. This indicates that the flooding gradient pose no overall effect on C_MAOM_ and C_oPOM_ but interacts with the salinity gradient and other factors leading to local variability. The OC in mineral association and aggregates occlusion at the salt marsh might be less affected by the flooding gradient due to generally smaller sedimentation rates, which also contain less clay (Butzeck et al., 2015). The negative effect of increasing flooding at the brackish marsh can be attributed to differences in the texture of the deposited sediments along the gradient that resulted in different soil textures. The brackish pioneer zone was characterized by high sand and very low clay contents, while the brackish low marsh had higher silt and clay contents with a larger potential for mineral association and aggregate formation (Table 1). However, high sand or clay contents were not always associated with low or high amounts of C_MAOM_ and C_oPOM_, such as in topsoil of the brackish high marsh and the freshwater marsh. Besides soil texture, the soil water content and dry-wet cycles are controlling aggregate dynamics and associated OM turnover in soil (Six et al., 2002b; Six et al., 2004; Najera et al., 2020). Aggregate formation and stability in soil can decrease with increasing flooding intensity and increasing soil water content in strongly flood-effected soils (Mao et al., 2018; Liu et al., 2021; Ran et al., 2021). This indicates that the conditions in the less inundated high marshes favor aggregation. In addition, the amounts of C_oPOM_ tended to be higher in topsoils compared to subsoils, which might also be related to less favorable conditions for aggregate formation in the predominantly wet subsoils. This is underlined by the observation that the amount of C_oPOM_ did not correlate with the clay content in subsoil. This supports the assumption that the conditions in less frequently flooded high marshes and topsoils promote the stabilization of OM in aggregates more effectively than the frequently inundated and overall wetter low marshes, pioneer zones and subsoils. Despite large aboveground biomass, high marshes did not show enriched C_fPOM_ pools. On the contrary, the contribution of C_fPOM_ to total SOC increased with increasing flooding at the salt and brackish marsh. This suggests that high marsh soils, due to their better aeration, are likely characterized by a faster turnover of plant litter, by which the OM is either lost by mineralization or is protected by mineral- association or in aggregates. The latter assumption is supported by the higher amount of C_oPOM_ and generally higher SOC contents in high marshes. In pioneer zones, the preservation of C_fPOM_ might benefit from the constant sedimentation and flooding, inducing oxygen-limitation and restricted mineralization (Schmidt et al., 2011). This is supported by a positive correlation between proportions of C_fPOM_ and the reduction index for the sampling month (RI_march_) while C_fPOM_ proportions were unrelated to the aboveground biomass.

### 4.3 Implications for climate change effects on SOC storage and stabilization in estuarine marshes

Various climatic- and non-climatic controls have led to a substantial loss in global wetland area. Sea-level rise and storm surges associated with climate change will pose a major impact on coastal ecosystems and their C storage capacity (IPCC, 2022). Landward expansion of marshes, vertical accretion, elevated atmospheric CO_2_, and increasing temperatures can enhance C sequestration in coastal ecosystems (Kirwan and Mudd, 2012; Macreadie et al., 2019; Lovelock and Reef, 2020). However, in our study we found decreasing SOC storage with increasing salinity and flooding along the estuarine gradients of the Elbe Estuary. This indicates the special role of estuaries at the aquatic-terrestrial interface. Particularly changes in SOC stabilization mechanisms should be considered when addressing implications of sea-level rise on C storage in these ecosystems. Submergence, burial, and erosion can pose a threat on coastal C storage if marshes cannot keep pace with sea-level rise (Macreadie et al., 2019; Lovelock and Reef, 2020; IPCC, 2022). The ability of landward migration is often limited by anthropogenic infrastructure (e.g. construction of dikes and weirs) (Lovelock and Reef, 2020; IPCC, 2022). In the Elbe Estuary this might not only limits the expansion of marshes but may also decrease SOC sequestration due to decreasing OC inputs and reduced SOC stabilization. Currently, SOC storage and stabilization in the freshwater marsh of the Elbe Estuary benefits from the strong fluvial influence, increasing biomass production, and the input of allochthonous OC and fine textured sediments. This beneficial fluvial influence might be reduced by sea-level rise and increased storm surges which enhance salinity and the contribution of coarse material in sediment deposits. Moreover, more frequent drought spells might reduce river discharge (IPCC, 2022), further reducing fine particle deposition and freshwater input. It was also shown that increasing inundation frequency and higher water content may have negative effects on aggregate formation and stability (Najera et al., 2020; Liu et al., 2021). Increasing dry-wet cycles can lead to the formation of macroaggregates at the expense of microaggregates with a strong implication for SOC turnover (Najera et al., 2020). How these effects control the SOC budget and stability in estuarine soils remains subject of future investigations.

## 5 Conclusion

In this study, we investigated effects of estuarine gradients (salinity and flooding) on SOC storage and potential stabilization mechanisms, indicated by SOC density fractions. Our results show that SOC stocks decrease from freshwater to salt marshes within the Elbe Estuary, which is likely related to decreasing autochthonous and allochthonous OC input downstream of the river. Mineral-association was the dominant stabilization mechanisms in the Elbe Estuary, followed by aggregation, reflected by a strong correlation between SOC stocks, MAOM, aggregate-occluded OC, and the clay content of the marsh soils. This indicates that the sedimentation dynamic is an important driver of SOC storage and stabilization mechanisms. Higher clay content led to higher potential of SOC preservation and stabilization at the freshwater marsh compared to salt and brackish sites. On the local scale, the actual degree of protection of SOC depends on the interaction of the estuarine gradients with each other and other local conditions; towards terrestrial influence (less flooding and lower salinity) the protection of SOC by aggregation increases. In contrast, free POM is more relevant for SOC storage in frequently-flooded parts of the marsh induced by reducing soil conditions.

Our data also highlights the strong spatial heterogeneity in SOC storage and stabilization, as well as environmental drivers within the estuary. Comparing our results with studies from other estuaries, indicated a strong variation in SOC storage and in the importance of controlling factors from one estuary to the other. Therefore, it is crucial to consider this heterogeneity in the assessment of SOC storage and stabilization to understand potential effects of climate change.

## Authors’ contributions

**Friederike Neiske:** Conceptualization, Formal Analysis, Investigation, Project administration, Visualization, Writing – original draft

**Maria Seedtke:** Investigation, Writing – review & editing

**Annette Eschenbach:** Funding acquisition, Resources, Supervision, Conceptualization, Writing – review & editing

**Monica Wilson:** Investigation, Writing – review & editing

**Kai Jensen:** Funding acquisition, Resources, Investigation, Supervision, Writing – review & editing

**Joscha N. Becker:** Conceptualization, Formal Analysis, Methodology, Project administration, Supervision, Writing – review & editing

## Acknowledgement

This study was funded by the Deutsche Forschungsgemeinschaft (DFG, German Research Foundation) within the Research Training Group 2530: “Biota-mediated effects on Carbon cycling in Estuaries” (project number 407270017; contribution to Universität Hamburg and Leibniz-Institut für Gewässerökologie und Binnenfischerei im Forschungsverbund Berlin e.V.). We thank Volker Kleinschmidt, Deborah Harms, Sumita Rui, Annika Naumann, Julian Mittmann-Goetsch, Fay Lexmond and involved student assistants for their help during field and lab work, as well as technical support.

## References

Abril G, Nogueira M, Etcheber H, Cabeçadas G, Lemaire E, Brogueira M. Behaviour of Organic Carbon in Nine Contrasting European Estuaries. Estuarine, coastal and shelf science 2002;54(2):241–62.

Barry A, Ooi SK, Helton AM, Steven B, Elphick CS, Lawrence BA. Carbon Dynamics Vary Among Tidal Marsh Plant Species in a Sea-level Rise Experiment. Wetlands 2023;43(7).

Boehlich MJ, Strotmann T. The Elbe Estuary. Die Küste 2008;74(1):288–306.

Butzeck C, Eschenbach A, Gröngröft A, Hansen K, Nolte S, Jensen K. Sediment Deposition and Accretion Rates in Tidal Marshes Are Highly Variable Along Estuarine Salinity and Flooding Gradients. Estuaries and Coasts 2015;38(2):434–50.

Carstens M, Claussen U, Bergemann M, Gaumert T. Transitional waters in Germany: the Elbe estuary as an example. Aquatic Conservation: Marine and Freshwater Ecosystems 2004;14(S1):S81–S92.

Castenson KL, Rabenhorst MC. Indicator of reduction in soil (IRIS) evaluation of a new approach for assessing reduced conditions in soil. Soil Science Soc of Amer J 2006;70(4):1222–6.

Chapman SK, Hayes MA, Kelly B, Langley JA. Exploring the oxygen sensitivity of wetland soil carbon mineralization. Biology letters 2019;15(1):20180407.

Chen C, Hall SJ, Coward E, Thompson A. Iron-mediated organic matter decomposition in humid soils can counteract protection. Nature communications 2020;11(1):2255.

Chmura GL, Anisfeld SC, Cahoon DR, Lynch JC. Global carbon sequestration in tidal, saline wetland soils. Global Biogeochemical Cycles 2003;17(4).

Craft C. Freshwater input structures soil properties, vertical accretion, and nutrient accumulation of Georgia and U.S tidal marshes. Limnology & Oceanography 2007;52(3):1220–30.

Craft C, Clough J, Ehman J, Joye S, Park R, Pennings S et al. Forecasting the effects of accelerated sea-level rise on tidal marsh ecosystem services. Frontiers in Ecol & Environ 2009;7(2):73–8.

Cui J, Li Z, Liu Z, Ge B, Fang C, Zhou C et al. Physical and chemical stabilization of soil organic carbon along a 500-year cultived soil chronosequence originating from estuarine wetlands: Temporal patterns and land use effects. Agriculture, Ecosystems & Environment 2014;196:10–20.

Drake K, Halifax H, Adamowicz SC, Craft C. Carbon Sequestration in Tidal Salt Marshes of the Northeast United States. Environmental management 2015;56(4):998–1008.

DWD. Time series and trends for the parameters temperature and precipitation: Reference period: 1991 - 2020. Federal State of Schleswig-Holstein, 2024. https://www.dwd.de/EN/ourservices/zeitreihen/zeitreihen.html#buehneTop (accessed February 20, 2024).

Engels JG, Jensen K. Patterns of wetland plant diversity along estuarine stress gradients of the Elbe (Germany) and Connecticut (USA) Rivers. Plant Ecology & Diversity 2009;2(3):301–11.

Enwright NM, Griffith KT, Osland MJ. Barriers to and opportunities for landward migration of coastal wetlands with sea-level rise. Frontiers in Ecol & Environ 2016;14(6):307–16.

Golchin A, Oades JM, Skjemstad JO, Clarke P. Study of free and occluded particulate organic matter in soils by solid state 13C Cp/MAS NMR spectroscopy and scanning electron microscopy. Soil Res. 1994;32(2):285.

Gorham C, Lavery P, Kelleway JJ, Salinas C, Serrano O. Soil Carbon Stocks Vary Across Geomorphic Settings in Australian Temperate Tidal Marsh Ecosystems. Ecosystems 2021;24(2):319–34.

Graves S, Piepho H-P, Selzer ML. Package ‘multcompView’. Visualizations of paired comparisons 2015.

Hansen K, Butzeck C, Eschenbach A, Gröngröft A, Jensen K, Pfeiffer E-M. Factors influencing the organic carbon pools in tidal marsh soils of the Elbe estuary (Germany). J Soils Sediments 2017;17(1):47–60.

Hothorn T, Bretz F, Westfall P, Heiberger RM, Schuetzenmeister A, Scheibe S et al. Package ‘multcomp’. Simultaneous inference in general parametric models. Project for Statistical Computing, Vienna, Austria 2016.

Human LRD, Els J, Wasserman J, Adams JB. Blue carbon and nutrient stocks in salt marsh and seagrass from an urban African estuary. The Science of the total environment 2022;842:156955.

IPCC, editor. Climate Change 2022 – Impacts, Adaptation and Vulnerability: Contribution of Working Group II to the Sixth Assessment Report of the Intergovernmental Panel on Climate Change. [H.-O. Pörtner, D.C. Roberts, M. Tignor, E.S. Poloczanska, K. Mintenbeck, A. Alegría, M. Craig, S. Langsdorf]. Cambridge, UK and New York, NY, USA: Cambridge University Press; 2022.

Kappenberg J, Grabemann I. Variability of the mixing zones and estuarine turbidity maxima in the Elbe and Weser estuaries. Estuaries 2001;24:699–706.

Kauffman JB, Giovanonni L, Kelly J, Dunstan N, Borde A, Diefenderfer H et al. Total ecosystem carbon stocks at the marine-terrestrial interface: Blue carbon of the Pacific Northwest Coast, United States. Global Change Biology 2020;26(10):5679–92.

Kelleway JJ, Saintilan N, Macreadie PI, Ralph PJ. Sedimentary Factors are Key Predictors of Carbon Storage in SE Australian Saltmarshes. Ecosystems 2016;19(5):865–80.

Kirwan ML, Megonigal JP. Tidal wetland stability in the face of human impacts and sea-level rise. Nature 2013;504(7478):53–60.

Kirwan ML, Mudd SM. Response of salt-marsh carbon accumulation to climate change. Nature 2012;489(7417):550–3.

Kleber M, Eusterhues K, Keiluweit M, Mikutta C, Mikutta R, Nico PS. Mineral–Organic Associations: Formation, Properties, and Relevance in Soil Environments. In: Elsevier; 2015. p. 1–140.

Liu Y, Ma M, Ran Y, Yi X, Wu S, Huang P. Disentangling the effects of edaphic and vegetational properties on soil aggregate stability in riparian zones along a gradient of flooding stress. Geoderma 2021;385:114883.

Lovelock CE, Reef R. Variable Impacts of Climate Change on Blue Carbon. One Earth 2020;3(2):195–211.

Lützow Mv, Kögel-Knabner I, Ekschmitt K, Matzner E, Guggenberger G, Marschner B, et al. Stabilization of organic matter in temperate soils: mechanisms and their relevance under different soil conditions – a review. European J Soil Science 2006;57(4):426–45.

Macreadie PI, Anton A, Raven JA, Beaumont N, Connolly RM, Friess DA et al. The future of Blue Carbon science. Nature communications 2019;10(1):3998.

Maietta CE, Bernstein ZA, Gaimaro JR, Buyer JS, Rabenhorst MC, Monsaint-Queeney VL et al. Aggregation but Not Organo-Metal Complexes Contributed to C Storage in Tidal Freshwater Wetland Soils. Soil Science Soc of Amer J 2019;83(1):252–65.

Mao R, Ye S-Y, Zhang X-H. Soil-Aggregate-Associated Organic Carbon Along Vegetation Zones in Tidal Salt Marshes in the Liaohe Delta. CLEAN Soil Air Water 2018;46(4).

Maxwell TL, Rovai AS, Adame MF, Adams JB, Álvarez-Rogel J, Austin WEN et al. Global dataset of soil organic carbon in tidal marshes. Scientific data 2023;10(1):797.

Mazarrasa I, Neto JM, Bouma TJ, Grandjean T, Garcia-Orellana J, Masqué P et al. Drivers of variability in Blue Carbon stocks and burial rates across European estuarine habitats. The Science of the total environment 2023;886:163957.

Mcleod E, Chmura GL, Bouillon S, Salm R, Björk M, Duarte CM et al. A blueprint for blue carbon: toward an improved understanding of the role of vegetated coastal habitats in sequestering CO 2. Frontiers in Ecol & Environ 2011;9(10):552–60.

Mittmann-Goetsch J, Wilson M, Jensen K, Mueller P. Wetland roots as soil reducers – Insights from a Wadden Sea salt-marsh study; 2024, Preprint.

Mueller P, Granse D, Nolte S, Weingartner M, Hoth S, Jensen K. Unrecognized controls on microbial functioning in Blue Carbon ecosystems: The role of mineral enzyme stabilization and allochthonous substrate supply. Ecology and evolution 2020;10(2):998–1011.

Müller H-W, Dohrmann R, Klosa D, Rehder S, Eckelmann W. Comparison of two procedures for particle-size analysis: Köhn pipette and X-ray granulometry. Z. Pflanzenernähr. Bodenk. 2009;172(2):172–9.

Najera F, Dippold MA, Boy J, Seguel O, Koester M, Stock S et al. Effects of drying/rewetting on soil aggregate dynamics and implications for organic matter turnover. Biol Fertil Soils 2020;56(7):893–905.

R Core Team. R. A language and environment for statistical computing. Vienna, Austria: R Foundation for Statistical Computing; 2022.

Rabenhorst MC. Protocol for using and interpreting IRIS tubes. Soil Survey Horizons 2008;49(3):74–7.

Ran Y, Ma M, Liu Y, Zhou Y, Sun X, Wu S et al. Hydrological stress regimes regulate effects of binding agents on soil aggregate stability in the riparian zones. CATENA 2021;196:104815.

Rocci KS, Lavallee JM, Stewart CE, Cotrufo MF. Soil organic carbon response to global environmental change depends on its distribution between mineral-associated and particulate organic matter: A meta-analysis. The Science of the total environment 2021;793:148569.

Ruiz-Fernández AC, Carnero-Bravo V, Sanchez-Cabeza JA, Pérez-Bernal LH, Amaya-Monterrosa OA, Bojórquez-Sánchez S et al. Carbon burial and storage in tropical salt marshes under the influence of sea level rise. The Science of the total environment 2018;630:1628–40.

Schmidt MWI, Torn MS, Abiven S, Dittmar T, Guggenberger G, Janssens IA et al. Persistence of soil organic matter as an ecosystem property. Nature 2011;478(7367):49–56.

Schulte Ostermann T, Kleyer M, Heuner M, Fuchs E, Temmerman S, Schoutens K et al. Hydrodynamics affect plant traits in estuarine ecotones with impact on carbon sequestration potentials. Estuarine, coastal and shelf science 2021;259:107464.

Seyfferth AL, Bothfeld F, Vargas R, Stuckey JW, Wang J, Kearns K et al. Spatial and temporal heterogeneity of geochemical controls on carbon cycling in a tidal salt marsh. Geochimica et Cosmochimica Acta 2020;282:1–18.

Six J, Bossuyt H, Degryze S, Denef K. A history of research on the link between (micro)aggregates, soil biota, and soil organic matter dynamics. Soil and Tillage Research 2004;79(1):7–31.

Six J, Conant RT, Paul EA, Paustian K. Stabilization mechanisms of soil organic matter: Implications for C-saturation of soils. Plant and Soil 2002a;241(2):155–76.

Six J, Feller C, Denef K, Ogle SM, Moraes JC de, Albrecht A. Soil organic matter, biota and aggregation in temperate and tropical soils - Effects of no-tillage. Agronomie 2002b;22(7-8):755–75.

Spohn M, Babka B, Giani L. Changes in soil organic matter quality during sea-influenced marsh soil development at the North Sea coast. CATENA 2013;107:110–7.

Spohn M, Giani L. Carbohydrates, carbon and nitrogen in soils of a marine and a brackish marsh as influenced by inundation frequency. Estuarine, coastal and shelf science 2012;107:89–96.

Sun H, Jiang J, Cui L, Feng W, Wang Y, Zhang J. Soil organic carbon stabilization mechanisms in a subtropical mangrove and salt marsh ecosystems. The Science of the total environment 2019;673:502–10.

Tang H, Nolte S, Jensen K, Rich R, Mittmann-Goetsch J, Mueller P. Warming accelerates belowground litter turnover in salt marshes – insights from a Tea Bag Index study. Biogeosciences 2023;20(10):1925–35.

van de Broek M, Temmerman S, Merckx R, Govers G. Controls on soil organic carbon stocks in tidal marshes along an estuarine salinity gradient. Biogeosciences 2016;13(24):6611–24.

van de Broek M, Vandendriessche C, Poppelmonde D, Merckx R, Temmerman S, Govers G. Long-term organic carbon sequestration in tidal marsh sediments is dominated by old-aged allochthonous inputs in a macrotidal estuary. Global Change Biology 2018;24(6):2498–512.

Viret F, Grand S. Combined Size and Density Fractionation of Soils for Investigations of Organo-Mineral Interactions. Journal of visualized experiments JoVE 2019(144).

Visser JM, Duke-Sylvester SM, Carter J, Broussard WP. A Computer Model to Forecast Wetland Vegetation Changes Resulting from Restoration and Protection in Coastal Louisiana. Journal of Coastal Research 2013;67:51–9.

Wickham H, Chang W, Wickham MH. Package ‘ggplot2’. Create elegant data visualisations using the grammar of graphics. Version 2016;2(1):1–189.

Więski K, Guo H, Craft CB, Pennings SC. Ecosystem Functions of Tidal Fresh, Brackish, and Salt Marshes on the Georgia Coast. Estuaries and Coasts 2010;33(1):161–9.

Wiesmeier M, Urbanski L, Hobley E, Lang B, Lützow M von, Marin-Spiotta E, et al. Soil organic carbon storage as a key function of soils - A review of drivers and indicators at various scales. Geoderma 2019;333:149–62.

Wong VNL, Greene RSB, Dalal RC, Murphy BW. Soil carbon dynamics in saline and sodic soils: a review. Soil Use and Management 2010;26(1):2–11.

World reference base for soil resources 2022: International soil classification system for naming soils and creating legends for soil maps. 4th ed. Vienna, Austria: International Union of Soil Sciences; 2022.

Yuan Y, Li X, Jiang J, Xue L, Craft CB. Distribution of organic carbon storage in different salt-marsh plant communities: A case study at the Yangtze Estuary. Estuarine, coastal and shelf science 2020;243:106900.

Zhou J, Wu Y, Kang Q, Zhang J. Spatial variations of carbon, nitrogen, phosphorous and sulfur in the salt marsh sediments of the Yangtze Estuary in China. Estuarine, coastal and shelf science 2007;71(1-2):47–59.

