## supplement for "Soil organic carbon stocks and stabilization mechanisms in tidal marshes along estuarine gradients"

### Supplementary Material

Table S 1: Plant species with area coverage [%] and Ellenberg Indicator Values for salinity (EIV-S) along the salinity and flooding gradient of the Elbe Estuary in July 2022.

| **Salt High Marsh** | [%] | **Brackish High Marsh** | [%] | **Fresh High Marsh** | [%] |
|---|---|---|---|---|---|
| *Agrostis stolonifera* | <0.1 | *Agrostis stolonifera* | 0.1 | *Agrostis stolonifera* | 0.8 |
| *Aster tripolium* | 13.4 | *Berula erecta* | 0.3 | *Calystegia sepium* | 7.2 |
| *Atriplex littoralis* | 0.2 | *Calystegia sepium* | 28.3 | *Equisetum palustre* | 0.1 |
| *Atriplex prostrata* | 15.8 | *Dactylis glomerata* | <0.1 | *Phalaris arundinacea* | 7.6 |
| *Elymus athericus* | 62.3 | *Epilobium hirsutum* | 24.1 | *Phragmites australis* | 54.0 |
| *Salicornia europaea* | <0.1 | *Galeopsis sp.* | <0.1 | *Urtica dioica* | 21.1 |
| *Suaeda maritima* | 0.5 | *Galium palustre* | 0.2 | *Valeriana officinalis* | 0.6 |
| | | *Phalaris arundinacea* | 4.9 | | |
| | | *Phragmites australis* | 10.9 | | |
| | | *Poa trivalis* | 0.2 | | |
| | | *Rosa sp.* | 2.3 | | |
| | | *Urtica dioica* | 6.7 | | |
| | | *Valeriana officinalis* | <0.1 | | |
| | | *Vicia cracca* | 0.3 | | |
| EIV-S: 5.4 | | EIV-S: 0.3 | | EIV-S: 0.0 | |
| **Salt Low Marsh** | [%] | **Brackish Low Marsh** | [%] | **Fresh Low Marsh** | [%] |
| *Agrostis stolonifera* | 0.2 | *Agrostis stolonifera* | 0.2 | *Bidens sp.* | 0.1 |
| *Aster tripolium* | 18.2 | *Berula erecta* | 6.9 | *Caltha palustris* | 0.1 |
| *Atriplex littoralis* | <0.1 | *Caltha palustris* | 0.3 | *Myosotis palustris* | 0.1 |
| *Atriplex prostrata* | 5.6 | *Calystegia sepium* | 9.5 | *Nasturtium officinale* | 0.1 |
| *Elymus athericus* | <0.1 | *Lycopus europaeus* | 0.9 | *Phragmites australis* | 92.6 |
| *Festuca rubra* | 8.3 | *Phragmites australis* | 73.0 | | |
| *Halimione portulacoides* | 1.4 | | | | |
| *Plantago maritima* | <0.1 | | | | |
| *Puccinellia maritima* | 40.8 | | | | |
| *Salicornia europaea* | 1.7 | | | | |
| *Spartina anglica* | 2.3 | | | | |
| *Spergularia media* | 0.1 | | | | |
| *Suaeda maritima* | 3.4 | | | | |
| *Triglochin maritima* | 13.2 | | | | |
| EIV-S: 7.0 | | EIV-S: 0.1 | | EIV-S: 0.0 | |
| **Salt Pioneer Zone** | [%] | **Brackish Pioneer Zone** | [%] | **Fresh Pioneer Zone** | [%] |
| *Aster tripolium* | 3.2 | *Bolboschoenus maritimus* | 73.5 | *Alisma plantago-aquatica* | 1.8 |
| *Atriplex prostrata* | 2.0 | *Nasturtium officinale* | 0.2 | *Caltha palustris* | 16.0 |
| *Halimione portulacoides* | 0.01 | | | *Mentha aquatica* | 0.1 |
| *Puccinellia maritima* | 4.1 | | | *Myosotis palustris* | 0.1 |
| *Salicornia europaea* | <0.1 | | | *Typha angustifolia* | 46.0 |
| *Spartina anglica* | 59.1 | | | | |
| *Suaeda maritima* | 2.6 | | | | |
| *Triglochin maritima* | 0.8 | | | | |
| EIV-S: 7.8 | | EIV-S: 2.0 | | EIV-S: 0.7 | |

Table S 2: SOC stocks (t $ha^{-1}$), SOC contents (%), amount of SOC fractions (mg C $g^{-1}$ soil) and proportions of SOC fractions (% of total SOC) ($C_{MAOM}$ = mineral-associated OC, $C_{oPOM}$ = aggregate-occluded OC, and $C_{fPOM}$ = OC in free particulate organic matter) of the different marsh types and zones along the salinity (salt marsh, brackish marsh, freshwater marsh) and flooding gradient (HM = high marsh, LM = low marsh, PZ = pioneer zone) of the Elbe Estuary (mean ± standard error, n = 5).

| Marsh type & zone | | Depth [cm] | SOC | | | $C_{MAOM}$ | | $C_{oPOM}$ | | $C_{fPOM}$ | |
|---|---|---|---|---|---|---|---|---|---|---|---|
| | | | stocks | stocks | contents | amount | proportion | amount | proportion | amount | proportion |
| Salt Marsh | HM | 0 - 10 | 28.7 ± 2.6 | 73.7 ± 4.9 | 2.9 ± 0.6 | 14.1 ± 1.7 | 48.8 ± 4.6 | 8.1 ± 2.3 | 27.3 ± 4.6 | 3.1 ± 1.1 | 11.0 ± 4.8 |
| | | 10 - 30 | 45.0 ± 2.6 | | 2.4 ± 0.3 | 17.1 ± 3.0 | 69.8 ± 7.5 | 4.6 ± 1.8 | 18.7 ± 6.9 | 1.8 ± 0.9 | 7.7 ± 4.3 |
| | LM | 0 - 10 | 14.6 ± 0.7 | 39.9 ± 2.9 | 1.6 ± 0.2 | 8.6 ± 1.4 | 54.7 ± 4.4 | 3.3 ± 1.0 | 20.8 ± 4.0 | 1.9 ± 0.4 | 12.3 ± 3.8 |
| | | 10 - 30 | 25.3 ± 2.8 | | 1.5 ± 0.4 | 8.8 ± 2.7 | 60.4 ± 12.8 | 2.6 ± 1.0 | 17.7 ± 3.2 | 2.1 ± 0.4 | 15.2 ± 4.8 |
| | PZ | 0 - 10 | 14.4 ± 0.9 | 46.7 ± 2.7 | 1.8 ± 0.3 | 9.9 ± 1.9 | 54.4 ± 6.5 | 3.9 ± 0.8 | 21.3 ± 2.5 | 3.8 ± 1.0 | 21.1 ± 5.1 |
| | | 10 - 30 | 32.3 ± 3.3 | | 1.6 ± 0.4 | 10.3 ± 2.6 | 62.8 ± 10.3 | 3.1 ± 1.1 | 19.4 ± 6.8 | 1.8 ± 0.7 | 10.4 ± 2.4 |
| Brackish Marsh | HM | 0 - 10 | 31.4 ± 6.2 | 68.0 ± 7.8 | 3.8 ± 1.0 | 21.9 ± 7.9 | 48.6 ± 5.0 | 17.8 ± 8.4 | 38.1 ± 6.1 | 3.0 ± 1.7 | 5.9 ± 1.1 |
| | | 10 - 30 | 36.6 ± 2.7 | | 1.6 ± 0.3 | 7.6 ± 2.5 | 46.0 ± 8.7 | 5.4 ± 1.4 | 32.7 ± 4.5 | 0.8 ± 0.3 | 5.1 ± 2.1 |
| | LM | 0 - 10 | 23.1 ± 1.5 | 55.9 ± 4.2 | 2.7 ± 0.4 | 16.4 ± 1.0 | 62.5 ± 6.8 | 8.5 ± 1.0 | 33.0 ± 8.6 | 4.4 ± 2.1 | 16.3 ± 6.5 |
| | | 10 - 30 | 32.8 ± 3.4 | | 1.3 ± 0.3 | 6.2 ± 0.9 | 49.1 ± 7.6 | 3.4 ± 1.3 | 25.6 ± 5.9 | 4.0 ± 1.4 | 29.8 ± 5.2 |
| | PZ | 0 - 10 | 2.9 ± 0.3 | 9.3 ± 0.6 | 0.2 ± 0.0 | 0.9 ± 0.2 | 41.6 ± 10.4 | 1.3 ± 0.4 | 57.0 ± 18.4 | 0.7 ± 0.2 | 30.5 ± 3.4 |
| | | 10 - 30 | 6.4 ± 0.6 | | 0.2 ± 0.1 | 1.3 ± 0.5 | 52.9 ± 17.4 | 0.5 ± 0.3 | 22.3 ± 9.9 | 0.8 ± 0.3 | 32.1 ± 10.2 |
| Freshwater Marsh | HM | 0 - 10 | 30.2 ± 1.5 | 70.5 ± 2.6 | 5.2 ± 0.6 | 35.2 ± 8.0 | 67.0 ± 8.7 | 14.3 ± 2.6 | 27.9 ± 6.8 | 2.3 ± 1.2 | 4.5 ± 2.4 |
| | | 10 - 30 | 40.3 ± 1.7 | | 2.8 ± 0.3 | 20.7 ± 2.4 | 74.1 ± 8.2 | 4.4 ± 1.8 | 15.6 ± 6.0 | 1.2 ± 0.4 | 4.4 ± 1.5 |
| | LM | 0 - 10 | 32.3 ± 1.9 | 74.6 ± 4.3 | 5.8 ± 0.7 | 36.7 ± 1.1 | 64.8 ± 10.8 | 12.5 ± 2.4 | 21.6 ± 2.4 | 11.2 ± 3.6 | 19.3 ± 4.6 |
| | | 10 - 30 | 42.3 ± 3.0 | | 2.5 ± 0.4 | 11.4 ± 1.7 | 46.4 ± 5.8 | 6.6 ± 1.5 | 26.8 ± 3.3 | 8.6 ± 2.7 | 34.6 ± 8.4 |
| | PZ | 0 - 10 | 22.3 ± 0.5 | 69.3 ± 2.9 | 4.6 ± 0.2 | 32.4 ± 2.1 | 69.7 ± 3.9 | 8.5 ± 2.2 | 18.3 ± 5.2 | 4.0 ± 0.4 | 8.7 ± 1.3 |
| | | 10 - 30 | 47.0 ± 2.8 | | 4.2 ± 0.5 | 35.9 ± 3.1 | 81.8 ± 2.9 | 5.2 ± 2.2 | 12.4 ± 5.1 | 4.0 ± 1.1 | 9.8 ± 3.8 |

Table S 3: Effects of estuarine gradients on SOC stocks (t $ha^{-1}$), SOC contents (%), amounts (mg C $g^{-1}$ soil) and proportions of SOC fractions (% of total SOC) ($C_{MAOM}$ = mineral-associated OC, $C_{oPOM}$ = aggregate-occluded OC, and $C_{fPOM}$ = OC in free particulate organic matter). P- and F-values are derived from ANOVA and asterisk indicate significant differences (* $p < 0.05$; ** $p < 0.005$; *** $p < 0.001$) (df = degree of freedom).

| Gradient | Depth [cm] | | SOC | | $C_{MAOM}$ | | $C_{oPOM}$ | | $C_{fPOM}$ | |
|---|---|---|---|---|---|---|---|---|---|---|
| | | | stock | content | amount | proportion | amount | proportion | amount | proportion |
| Salinity | 0 - 10 | p | < 0.001 | < 0.001 | < 0.001 | < 0.001 | < 0.001 | < 0.001 | < 0.001 | < 0.001 |
| | | F | 13.6 | 188.2 | 166.7 | 22.9 | 16.0 | 30.9 | 16.8 | 12.4 |
| | | df | 2 | 2 | 2 | 2 | 2 | 2 | 2 | 2 |
| | 10 - 30 | p | < 0.001 | < 0.001 | < 0.001 | < 0.001 | < 0.001 | < 0.001 | < 0.001 | < 0.001 |
| | | F | 33.8 | 142.1 | 215.3 | 12.7 | 10.7 | 9.7 | 27.0 | 27.0 |
| | | df | 2 | 2 | 2 | 2 | 2 | 2 | 2 | 2 |
| Flooding | 0 - 10 | p | < 0.001 | < 0.001 | < 0.001 | 0.061 . | < 0.001 | 0.04 * | < 0.001 | < 0.001 |
| | | F | 35.9 | 42.5 | 21.6 | 3.0 | 28.1 | 3.4 | 17.2 | 35.7 |
| | | df | 2 | 2 | 2 | 2 | 2 | 2 | 2 | 2 |
| | 10 - 30 | p | < 0.001 | < 0.001 | < 0.001 | 0.002 ** | 0.006 ** | 0.050 . | < 0.001 | < 0.001 |
| | | F | 15.4 | 9.7 | 41.1 | 7.4 | 5.9 | 3.3 | 38.4 | 38.4 |
| | | df | 2 | 2 | 2 | 2 | 2 | 2 | 2 | 2 |
| Salinity: Flooding | 0 - 10 | p | < 0.001 | < 0.001 | < 0.001 | 0.004 | < 0.001 | < 0.001 | < 0.001 | < 0.001 |
| | | F | 7.6 | 17.3 | 10.2 | 4.7 | 5.9 | 6.0 | 16.1 | 16.1 |
| | | df | 4 | 4 | 4 | 4 | 4 | 4 | 4 | 4 |
| | 10 - 30 | p | < 0.001 | < 0.001 | < 0.001 | 0.002 ** | < 0.001 | 0.007 ** | < 0.001 | < 0.001 |
| | | F | 19.0 | 30.1 | 63.5 | 5.3 | 6.5 | 4.2 | 12.1 | 12.1 |
| | | df | 4 | 4 | 4 | 4 | 4 | 4 | 4 | 4 |
